# Somatic mutation detection from non-malignant liver tissues using cell-free DNA

**DOI:** 10.64898/2026.11.23.767883

**Authors:** Boyuan Li, Srinivas Rajagopalan, Alexandre Pellan Cheng, Min Zhu, Ashwini Arvind, Andrew Shubin, Mengmeng Wang, Julia Terrell, Lisa Quirk, Parsia A Vagefi, Chul Ahn, Roy Wang, Courtney Katz, Hiroaki Kanzaki, Fatma Kalkan, Joseph Gunderson, Jeon Lee, Natalia Brzozowska, Matthew Hoare, Peter J Campbell, John Zinno, William Driscoll, Adam Yopp, Yujin Hoshida, Tao Wang, Stephen Chung, David Hsieh, Amit Singal, Madhukar Patel, Dan A. Landau, Hao Zhu

**Author notes:** Co-corresponding: Correspondence and lead contact: Hao Zhu and Dan Landau. Equal contribution (Boyuan Li, Srinivas Rajagopalan, Alexandre Pellan Cheng).

## Abstract

Somatic mutations accumulate in the liver during aging, yet their role in chronic liver disease (CLD) remains unclear due to the difficulty of obtaining liver tissue for analysis. Here, we demonstrate that circulating cell-free DNA (cfDNA) can be used as a minimally invasive tool to detect liver-derived somatic mutations in patients with CLD. Using ultra-deep targeted duplex sequencing, we show that CLD patients have a higher mutation burden in CLD-associated somatic mutation (CASM) genes compared to healthy individuals. Recurrent driver mutations, including *CHD4*, *INSR*, and *GPAM*, were detected in both liver tissue and cfDNA, with distinct mutational patterns linked to disease etiology: *INSR* mutations were enriched in metabolic-dysfunction associated steatotic liver disease, while *GPAM* mutations were associated with alcohol-related liver disease. Liver transplantation led to a significant reduction in CASM mutation burden in cfDNA, providing strong evidence for the diseased-liver origin of these mutations. These findings establish cfDNA analysis as a scalable and noninvasive platform for investigating somatic evolution in non-malignant tissues, offering new opportunities for molecular stratification, longitudinal monitoring, and early detection of CLD progression.

## Introduction

The accumulation of genetic mutations in subpopulations of cells is an inevitable part of human aging, even in physiologically normal tissues^1–8^. Deep-sequencing studies over the past decade have shown that healthy organs are mosaic, with mutant clones that expand under positive selection rather than drift. A central theme is that many of these expanding clones carry fitness promoting mutations yet remain non-malignant, indicating that clonal expansion and carcinogenesis are distinct processes shaped by tissue-specific selective pressures and environmental exposures^9–15^. The fields biggest open questions are which clones actually progress to disease versus which are neutral or even protective, and whether this clonal evolution represents an underappreciated, universal driver of aging and age-related cancer risk.

Progress in this field has been constrained by the limited availability of non-malignant tissues, which has made it difficult to ascertain mutational landscapes on the population level^4,6,16^. In contrast to cancer genomics, in which samples are obtained routinely during surgery, non-malignant tissues are not routinely sampled. While recent work from Lawson et al. demonstrates that cheek swabs can be sequenced at epidemiological scale, this is impractical for most visceral organs^17^. The inability to define the mutational landscape of internal organs at scale has constrained the ability to advance the field of somatic genomics outside of clonal hematopoiesis. Establishing non-invasive assays for mutation detection would transform the field. First, this would enable population level studies to identify associations between somatic mutations and clinical variables. Second, this would allow longitudinal tracking of mutant clone dynamics during the course of disease, before and after clinical interventions. Third, it could provide a premalignant detection window to capture malignant transformation events. Circulating cfDNA is an attractive, minimally invasive alternative option for genomic analysis^18–20^. Although cfDNA testing has revolutionized cancer screening and prenatal diagnosis^21–23^, its application to non-malignant somatic mutations presents substantial challenges^24–26^. Among total cfDNA, mutation-bearing fragments derived from individual organs are expected to be extremely rare^27,28^. Their sensitive detection therefore requires methods with exceptionally high accuracy. Duplex sequencing provides such accuracy by independently validating variants on both strands of a DNA molecule^29,30^, and could be well suited for identifying rare somatic mutations at extremely low variant allele frequencies (VAFs)^31–33^. Even with accurate sequencing, the source organ for cfDNA is often unknown.

Despite these challenges, this approach could be more promising for the liver than for most organs. Hepatocytes are a major contributor to the circulating cfDNA pool, so liver-derived DNA is abundant in the blood. Second, dying cells shed their DNA into circulation, and CLD is defined by ongoing injury and cell death. Third, mutant clones within the liver can expand to grossly visible nodules that represent appreciable VAFs within the cfDNA compartment. Recent studies have shown that somatic mutations are the rule, rather than the exception, in CLD tissues^34–37^. Chronic liver injury is associated with clonal expansion of somatic mutations, which is particularly evident in the settings of metabolic dysfunction-associated steatotic liver disease (MASLD), alcohol-related liver disease (ALD), and chronic viral hepatitis^35,38,39^. While somatic evolution is widespread in CLD, the clinical significance of these mutations and their etiology-specific features remain poorly understood, in part because an insufficient number of patients have been sampled to know if these somatic mutations are associated with clinical features of disease.

Here, we performed targeted duplex sequencing of cfDNA to analyze somatic mutations in CLD. Across 303 samples from 224 patients, including pre-transplant cfDNA, post-transplant cfDNA, and liver tissue, we detected an increased mutational burden of liver-specific driver genes in the cfDNA of patients with CLD, and this mutational burden decreased significantly after liver transplantation. Comparative analyses revealed distinct mutational features among patients with different liver disease etiologies. These findings highlight the potential of cfDNA-based approaches for detecting non-neoplastic somatic mutations prior to cancer.

## Results

### Identification of driver gene mutations in liver tissues across different etiologies

To define the mutational landscape of CLD through cfDNA analysis, we assembled both liver tissue and plasma cohorts from patients with non-malignant liver disease (**Fig. 1a** and **Supplementary Table 1**). To establish a reference map of recurrent somatic mutations in diseased liver tissue, we first collected liver samples from 31 patients with diverse CLD etiologies, including 13 with ALD and 11 with MASLD (**Fig. 1b,c**). To investigate circulating cfDNA, we analyzed plasma samples from a cohort of 224 individuals, comprising a total of 272 samples. Among these, 216 samples were collected from 168 patients with CLD, alongside 41 healthy controls, and 15 disease controls with cardiac disease. Among patients with CLD, most had cirrhosis, defined as F4 fibrosis (n = 140); ALD (n = 53) and MASLD (n = 43) represented the largest etiologic groups. To assess the tissue origin of cfDNA-detected mutations, we also collected 53 post-transplant plasma samples from a subset of these patients, including one patient sampled after each of two separate liver transplantations. (**Fig. 1b,c**).

**Fig. 1:**
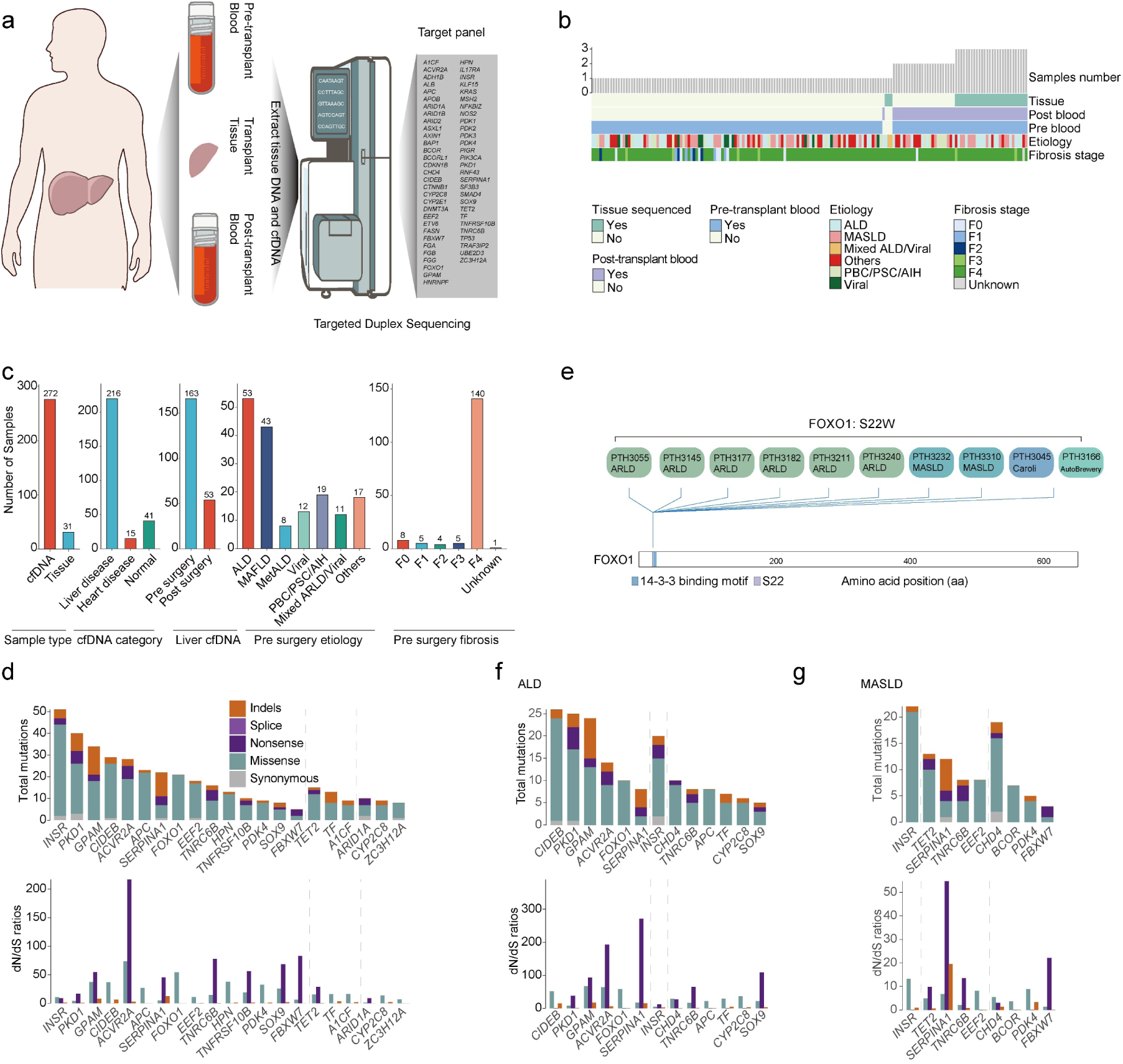
Study design, sample composition, and selection analyses in CLD liver tissues. a. Schematic overview of the study workflow. Pre-transplant blood, liver tissue, and post-transplant blood samples were collected, and tissue DNA and cfDNA were extracted and subjected to targeted duplex sequencing. The target gene panel used in this study is shown on the right. b. Overview of sample composition. The bar plot at the top shows the number of samples available for each subject. The annotation tracks below indicate the presence or absence of tissue samples, the presence or absence of pre/post-transplant blood samples, etiological group, and fibrosis stage. c. Summary of sample distribution. From left to right, the panels show the numbers of cfDNA and tissue samples, numbers of samples in different clinical categories (liver disease, heart disease controls, and normal controls), numbers of pre- and post-transplant samples in liver cfDNA, numbers of samples in each etiological group in pre surgery samples, and numbers of samples in each fibrosis stage category in pre surgery samples. Numbers above bars indicate sample counts. d. Gene-level selection analysis in the full tissue cohort. The upper panel shows the total number of mutations detected in each gene, stratified by mutation class into synonymous, missense, nonsense, splice and indel mutations; the lower panel shows the corresponding dN/dS ratios. Dashed lines separate genes identified at different significance thresholds. P values and q values were derived from dNdScv. Vertical dashed lines divide genes into three groups according to the statistical significance thresholds: genes to the left of the first dashed line meet *q* ≤ 0.05 and *p* ≤ 0.05; genes between the two dashed lines meet *q* ≤ 0.1 and *p* ≤ 0.05; genes to the right of the second dashed line have *q* > 0.1 but *p* ≤ 0.05. P values and q values were derived from dNdScv. e. Mutation spectrum of FOXO1 S22W in tissue. Lollipop plot showing the distribution of the recurrent FOXO1 S22W mutation detected in tissues from multiple individuals across disease etiologies. Each bubble represents one sample harboring the mutation, labeled by patient ID and disease category. The mutation maps to S22 of FOXO1 and lies adjacent to the annotated 14-3-3 binding motif. f. Gene-level selection analysis in tissue from the ALD subgroup. The upper panel shows the total number of mutations in each gene, stratified by mutation class, and the lower panel shows the corresponding dN/dS ratios. Vertical dashed lines divide genes into three groups according to the statistical significance thresholds: genes to the left of the first dashed line meet *q* ≤ 0.05 and *p* ≤ 0.05; genes between the two dashed lines meet *q* ≤ 0.1 and *p* ≤ 0.05; genes to the right of the second dashed line have *q* > 0.1 but *p* ≤ 0.05. P values and q values were derived from dNdScv. g. Gene-level selection analysis in tissue from the MASLD subgroup. The upper panel shows the total number of mutations in each gene, stratified by mutation class, and the lower panel shows the corresponding dN/dS ratios. Vertical dashed lines divide genes into three groups according to the statistical significance thresholds: genes to the left of the first dashed line meet *q* ≤ 0.05 and *p* ≤ 0.05; genes between the two dashed lines meet *q* ≤ 0.1 and *p* ≤ 0.05; genes to the right of the second dashed line have *q* > 0.1 but *p* ≤ 0.05. P values and q values were derived from dNdScv

All cfDNA and tissue samples were analyzed using a 59-gene targeted panel combined with duplex sequencing (**Extended Data Fig. 1a**). The panel included chronic liver disease (CLD)-associated somatic mutation (CASM) genes, liver lineage defining genes, hepatocellular carcinoma (HCC) genes, together with control gene sets comprising clonal hematopoiesis (CH) genes, non-disease control genes, colorectal cancer driver genes, and inflammatory bowel disease (IBD) genes^40–44^. Across the entire dataset, the median raw sequencing depth was 34,470x, with median single-strand consensus sequence (SSCs) depth of 6,690x and duplex consensus sequence (DCs) depth of 1,770x (**Extended Data Fig. 1b**). This study design enabled integrated analysis of somatic mutations across liver tissue, pre-transplant cfDNA, and post-transplant cfDNA, providing the necessary comparison samples to define liver-derived mutational profiles in CLD. To first define recurrent mutation events that are under evolutionary selection in our cohorts of non-malignant CLD tissues, we performed nonsynonymous/synonymous substitution (dN/dS) analysis^17,45^ on 30 diseased liver tissue samples, excluding one retransplant specimen. This dN/dS approach compares the rate of nonsynonymous substitutions, which alter protein sequences, with the rate of synonymous substitutions, which do not. Synonymous mutations are assumed to be selectively neutral and are used to estimate the background mutation rate, allowing detection of positive selection acting on protein-coding genes. A ratio of 1 indicates neutral evolution, <1 indicates negative (purifying) selection, and >1 indicates positive selection. To comprehensively evaluate the dN/dS results, we applied three significance thresholds: P value ≤ 0.05, FDR ≤ 0.1, and FDR ≤ 0.01.

This analysis identified genes previously reported as positively selected in CLD, including *PKD1, INSR, GPAM, CIDEB, FOXO1, ACVR2A,* and *TNRC6B*^34,37,46^ (**Fig. 1d**). In addition to these genes, we also identified *SERPINA1*. Although germline *SERPINA1* variants are well-established drivers of alpha-1 antitrypsin deficiency (AATD)-associated liver disease^47^, several truncating *SERPINA1* mutations were identified in our cohort. Recent work suggests that somatic truncating *SERPINA1* mutations can be positively selected in the liver as adaptive escape variants that reduce the toxic accumulation of mutant AAT protein^48^. In addition, *SOX9*, which has been implicated as an important gene in transcriptomic and protein-protein interaction network analyses of liver disease, also showed evidence of selection in our cohort^49,50^. Beyond these established genes, we identified *APC, EEF2, TNFRSF10B, PDK4* and *FBXW7* as previously unrecognized positively selected genes. Notably, our data also identified the previously reported *FOXO1*^S22W^ mutation^37^ (**Fig. 1e**), providing independent support for selective pressure acting on this hotspot mutation in CLD.

Selection of these driver gene candidates is biologically plausible in liver disease. Consistent with our identification of *INSR* as a positively selected gene, the liver is a major target organ of insulin signaling, and the INSR-IRS-PI3K-AKT axis is central to the regulation of gluconeogenesis, glycogen synthesis, and lipogenesis^51–53^. Prior functional studies have shown that impaired hepatic INSR signaling can protect against steatosis despite hyperinsulinemia and systemic insulin resistance, suggesting that clonal loss of *INSR* could confer a selective advantage in the chronically injured liver^54–57^. Likewise, decreased expression of the canonical tumor suppressor gene *APC* increases WNT signaling, raising the possibility that loss of *APC* may facilitate hepatocyte regeneration^58–60^. TNFRSF10B, a death receptor for TRAIL (TNFSF10), participates in apoptotic clearance of virus-infected cells^61,62^. *PDK4*, a key inhibitor of the mitochondrial pyruvate dehydrogenase complex, is upregulated in human MASH liver tissue and mouse MASH models and has been associated with lipid accumulation and insulin resistance^63–66^. Similarly, hepatocyte loss of *FBXW7* disrupts the ERRα-PPARα metabolic axis, suppresses fatty acid oxidation, and promotes endoplasmic reticulum stress, inflammation, and fibrosis, thereby accelerating MASH progression^67^. The positive selection of *FBXW7* observed here suggests that loss of this gene may enhance cellular fitness in the setting of liver injury. Additional genes meeting nominal selection criteria but supported by low mutation counts, such as *CDKN1B*, were considered exploratory and are reported in **Supplementary Table 2.**

Because CLD arises from distinct etiologies, we next asked whether different disease contexts impose distinct selective pressures on somatic mutations. Stratified selection analysis revealed clear etiology-specific patterns. In ALD, *GPAM* showed stronger evidence of positive selection, whereas in MASLD, mutations in *INSR*, *CHD4*, and *TNRC6B* were more strongly selected (**Fig. 1f,g**). These findings suggest that different etiologic contexts are associated with distinct selective environments, ultimately increasing the frequency of different driver mutations. To determine if these genes were differentially selected in specific CLD etiologies in independent patient cohorts, we analyzed a previously published dataset^37^ examining somatic mutations in MASLD and ALD. This analysis also showed that *GPAM* mutations were more positively selected in ALD, whereas *INSR*, *CHD4*, and *TNRC6B* were more strongly selected in MASLD (**Extended Data Fig. 1c,d**). Together, these findings demonstrate that distinct CLD etiologies are associated with different landscapes of somatic mutations that reflect the underlying selective pressures.

### Liver transplantation reveals the hepatic origin of cfDNA mutations

To determine if cfDNA can be used to characterize CASMs, we analyzed plasma samples from liver transplant recipients. Liver transplantation provides a natural experiment to test the tissue origin of cfDNA mutations, since the replacement of the native liver with a donor graft should specifically alter the liver-derived component of circulating cfDNA. We therefore compared pre- and post-transplant cfDNA samples from 53 patients, hypothesizing that transplantation would reshape cfDNA abundance and attenuate CASM mutational profiles in the plasma.

To quantify the liver-derived cfDNA fraction after transplantation, we leveraged donor-specific germline variants absent from the recipient (**Extended Data Fig. 2a,b**). To minimize any contribution of residual cfDNA released from the recipients native liver, we analyzed blood samples collected at least 22 days after liver transplantation. The median liver-derived cfDNA fraction was 2.5%, with a range of 0.4-12.0% across patients (**Fig. 2a**). These estimates are broadly consistent with previous reports describing organ-derived cfDNA fractions in the range of 0.4–2%^28,68,69^. These data indicate that the cfDNA fraction from the liver can be directly measured and varies substantially between individuals.

**Fig. 2:**
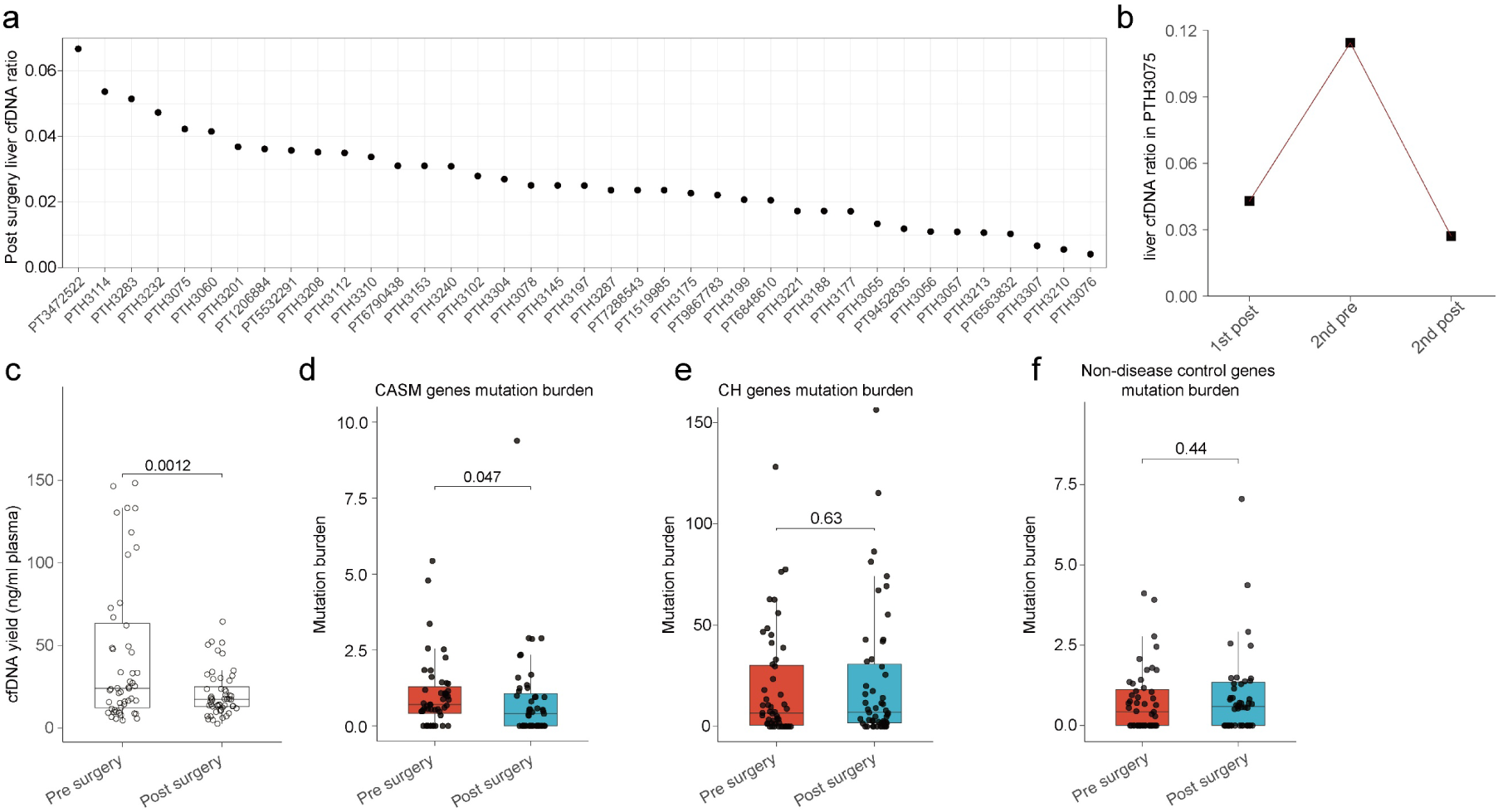
Estimation of the liver-derived cfDNA fraction after liver transplantation and changes in mutation burden. a. Estimated liver-derived cfDNA fraction in post-transplant samples. The y axis shows the liver cfDNA ratio in post-transplant samples, and the x axis shows individual post-transplant sample IDs, ordered by estimated fraction, from highest to lowest. b. Dynamic changes in the liver-derived cfDNA fraction in one patient (PTH3075) who underwent two independent liver transplantations. The y axis shows the liver cfDNA ratio, and the x axis shows different time points. Clear changes are observed after the first transplantation, before the second transplantation, and after the second transplantation. c. Comparison of plasma cfDNA yield before and after liver transplantation. The y axis shows cfDNA yield (ng/ml plasma), and the x axis shows the pre surgery and post surgery groups. P values for group comparisons are indicated. d. Comparison of mutation burden in CASM genes before and after liver transplantation. The y axis shows mutation burden, and the x axis shows the pre surgery and post surgery groups. P values for group comparisons are indicated. e. Comparison of mutation burden in CH genes before and after liver transplantation. The y axis shows mutation burden, and the x axis shows the pre surgery and post surgery groups. P values for group comparisons are indicated. f. Comparison of mutation burden in non-disease control genes before and after liver transplantation. The y axis shows mutation burden, and the x axis shows the pre surgery and post surgery groups. P values for group comparisons are indicated.

One patient underwent two independent liver transplantations, providing an opportunity to track liver-derived cfDNA dynamically across sequential graft replacement. In this individual, the estimated liver-derived cfDNA fraction was 4.3% after the first transplant, increased to 11.4% before the second transplant, and fell to 2.7% after the second transplant (**Fig. 2b**). The marked elevation observed before the second transplant, followed by a decline after graft replacement, suggests that liver-derived cfDNA can increase in the setting of a failing liver graft and provides a reference estimate for the upper range of liver-derived cfDNA contribution prior to transplantation.

We next compared pre-transplant cfDNA, explanted liver tissue, and post-transplant cfDNA to determine whether removal of the diseased native liver altered the mutational landscape ascertained in plasma. Following transplantation, the total plasma cfDNA abundance was significantly reduced (**Fig. 2c**), with median concentrations decreasing from 24.0 ng/ml pre-transplantation to 17.3 ng/ml post-transplantation, representing a 39% reduction. More importantly, mutation burden in CASM genes declined after transplantation, with the median burden decreasing from 0.7 mutations per million SSC molecules before transplantation to 0.4 mutations per million SSC molecules after transplantation, whereas mutation burden in CH genes and non-disease control genes remained largely unchanged (**Fig. 2d–f**). Thus, transplantation selectively reduced the CASM component of plasma cfDNA without broadly suppressing non-liver-associated mutational signals. Collectively, these findings provide strong evidence that the cirrhosis-associated mutational signal detected in cfDNA originates predominantly from the diseased native liver.

### cfDNA identifies liver disease-associated mutations and fragmentation patterns

To further characterize CASM detection with cfDNA in CLD, we compared plasma samples from patients with CLD and healthy controls. Healthy controls were individuals without known CLD or other major systemic illnesses. We first examined whether plasma cfDNA abundance differed between healthy individuals and patients with CLD. Those with CLD exhibited elevated plasma cfDNA concentrations relative to normal controls, with median levels of 8.6 ng/ml in normal samples, 12.6 ng/ml in F0–F3 samples, and 16.6 ng/ml in F4 samples. This corresponded to a 47% increase in F0–F3 and a 94% increase in F4 relative to normal controls (**Fig. 3a**), supporting the notion that the diseased liver contributes substantially to the cfDNA pool. Plasma cfDNA abundance increased progressively with cirrhosis decompensation as measured by the Child-Pugh score, a clinical scoring system used to assess the severity of CLD. The median levels of cfDNA were 13.1 ng/ml in Child–Pugh A, 16.1 ng/ml in Child-Pugh B and 22.2 ng/ml in Child–Pugh C, with the latter being significantly higher than both Child-Pugh A and B (**Fig. 3b**). This increasing cfDNA abundance is consistent with increased enhanced cell death, chronic inflammation, and fibrosis in end-stage liver disease^70–73^.

**Fig. 3:**
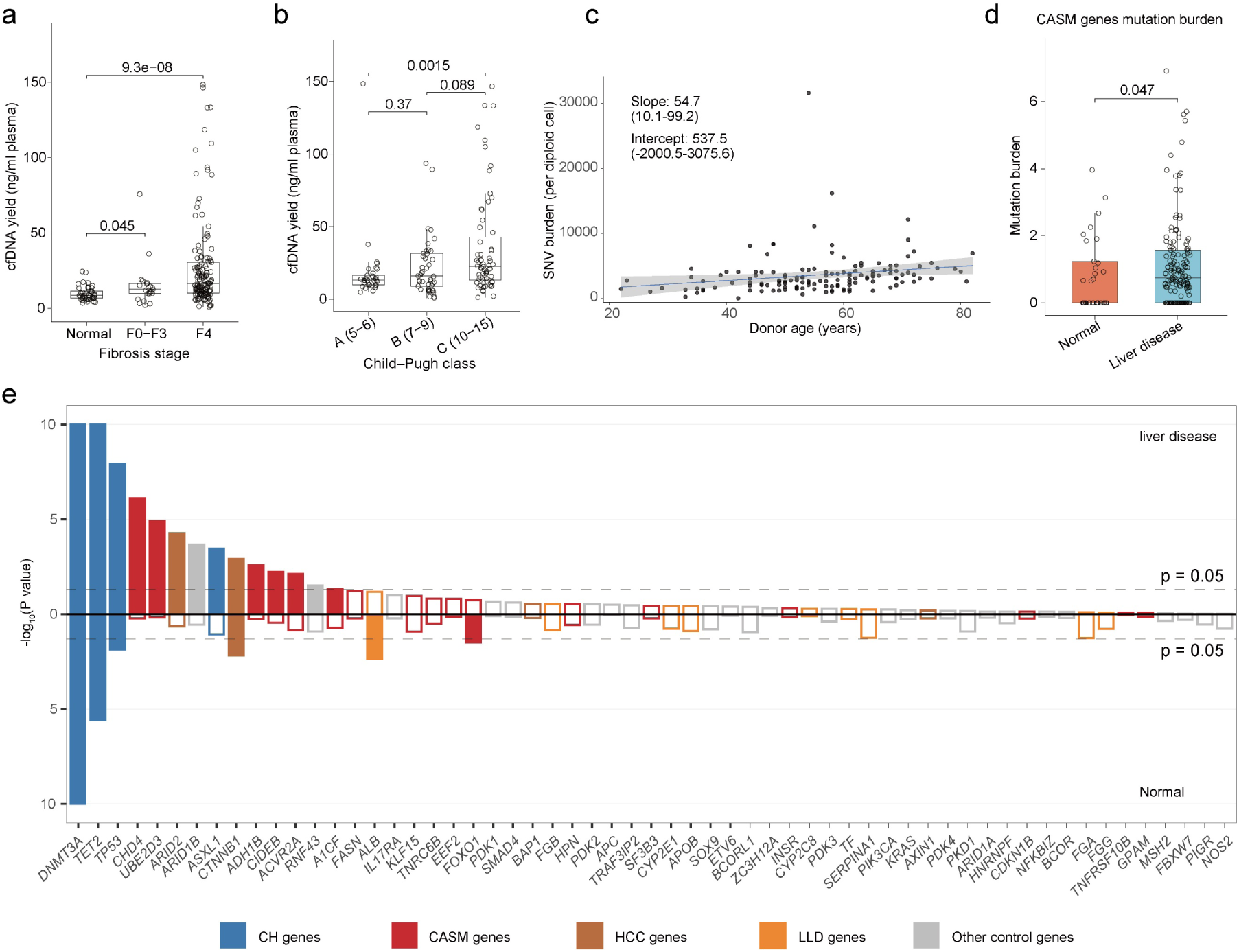
cfDNA yield, cirrhosis-associated driver gene mutation burden, and donor age–related analyses. a. Comparison of plasma cfDNA yield across fibrosis stage groups. The y axis shows cfDNA yield (ng/ml plasma), and the x axis shows the Normal, F0–F3, and F4 groups. P values for group comparisons are indicated. b. Comparison of plasma cfDNA yield across histological activity grades. The y axis shows cfDNA yield (ng/ml plasma), and the x axis shows groups A (5–6), B (7–9), and C (10–15). P values for group comparisons are indicated. c. Correlation analysis between donor age and SNV burden. The x axis shows donor age, and the y axis shows SNV burden (per diploid cell). The solid line indicates the linear fit, and the shaded area indicates the 95% confidence interval. The estimated slope and intercept, together with their corresponding intervals, are shown in the panel. d. Comparison of mutation burden in CASM genes. The y axis shows mutation burden, and the x axis shows the Normal, and liver disease (F0-F4) groups. P values for group comparisons are indicated. e. Selection analysis comparing liver disease samples (F0–F4) with normal controls at the gene level. The y axis shows −**log_10_**(***P***-value), where P values were calculated using the dNdScv package^45^, and the x axis shows gene names. Upward and downward bars represent liver disease and normal groups, respectively. The dashed lines indicate the significance threshold (P = 0.05), and non-significant genes are shown as unfilled bars. Different colors denote different gene categories, including CH genes, CASM genes, HCC genes, liver lineage defining (LLD) genes and other control genes.

Diseased liver tissues are characterized by elevated burdens of clonal somatic mutations, but these mutations are often present at low VAFs^35,36^. Because cfDNA released from individual mutant clones is diluted by DNA derived from the broader liver and other tissues, liver-derived somatic mutations are expected to occur at very low frequencies in plasma. To maximize sensitivity and accuracy for rare-variant detection, we performed ultra-deep targeted duplex sequencing of cfDNA. Across the 59-gene panel, we detected an average of 6.55 mutations per sample, including 1.54 mutations per sample in CASM genes (**Extended Data Fig. 3a**). Notably, individuals with liver disease harbored a higher burden of mutations in CASM genes than normal controls.

Aging and extent of CLD are both correlated with increased clonal expansion events identified in the liver. We asked if the same correlations could be observed in cfDNA from CLD patients. In cfDNA, we observed a trend toward increasing SNV burden as a function of age in patients with CLD. This age-associated increase was evident in cfDNA, with an estimated increase of approximately 55 SNVs per year, suggesting that the known accumulation of somatic mutation burden with age can be detected in cfDNA (**Fig. 3c**). Because CASMs are expected to arise from relatively small clones and occur at very low VAF in cfDNA, a critical benchmark for liver-derived signal is whether mutation burden is elevated in CLD patients relative to unaffected individuals. Consistent with this, mutation burden in CASM genes tended to be higher in patients with liver disease than in normal controls, with a median of 0 mutations per million SSC molecules in normal controls and 0.75 mutations per million SSC molecules in F4 samples (**Fig. 3d**). In contrast, no increase was observed in mutation burden for canonical CH or non-disease control genes (**Extended Data Fig. 3b,c**). These findings suggest that disease-associated genomic alterations in the liver are reflected in the plasma cfDNA mutational landscape.

To determine whether cfDNA could also recover CASM genes identified in tissue, we performed dN/dS analysis in cfDNA from CLD and non-CLD control patients. In controls, significant positive selection was limited to canonical CH genes including *DNMT3A, TET2,* and *TP53*^42,74^, together with *CTNNB1* and *FOXO1* (**Fig. 3e** and **Extended Data Fig. 3d,e**). In contrast, cfDNA from CLD patients revealed significant selection in CASM genes, including *CHD4*, *UBE2D3*, *AH1B*, *CIDEB*, and *ACVR2A*. Notably, *CHD4*, *CIDEB*, and *ACVR2A* were also positively selected in diseased liver tissue. Overall, CASM genes showed stronger selection signals in patient cfDNA than in normal controls, supporting the idea that cfDNA can recover driver mutational signals similar to those observed in tissue.

We next examined whether CLD was associated with altered cfDNA fragmentation patterns. Although previous work has characterized cfDNA fragmentomic features^75^, end-motif features have not yet been explored. Here, we specifically investigated the end-motif landscape of cfDNA in CLD. cfDNA from patients with cirrhosis showed a 4-bp reduction in median fragment length compared with normal controls, with the most pronounced depletion observed among longer cfDNA fragments (**Extended Data Fig. 3f**). In addition, cirrhosis samples showed altered end-motif composition compared with normal controls (**Extended Data Fig. 3g**). Further decomposition of fragmentation features demonstrated significant reductions in the DNASE1L3-associated F-profile I and the DNASE1-associated F-profile II, together with increases in F-profile IV and V in cirrhosis (**Extended Data Fig. 3h,i**). These alterations in end-motif profiles were fully or partially reversed following liver transplantation (**Extended Data Fig. 3j,k**). These changes could reflect attenuation of the dominant nuclease signatures in circulating blood, increased contribution of cfDNA from diseased liver, or altered patterns of cell death^76–81^. Notably, clustering based on fragmentation features separated liver disease samples from non-diseased controls, suggesting that cfDNA fragmentation could serve as an orthogonal biomarker of CLD. Together, these results show that even in non-malignant CLD, cfDNA differs markedly from normal plasma at multiple levels, including total abundance, somatic mutation burden, driver gene selection, fragment size, and end-fragmentation features.

### cfDNA mutations are concordant with liver-derived mutations

To further support the hypothesis that somatic mutations detected in plasma cfDNA reflect liver-derived alterations, we sought to identify previously reported CASMs identified from liver tissues. Given that Ng et al. identified *FOXO1^S22W^* as a hotspot mutation in steatotic liver disease^37^, its detection in cfDNA strongly suggests the ability to capture somatic mutations originating from the liver. In our cohort, *FOXO1^S22W^* was detected in the cfDNA from six pre-transplant liver disease patients, one post-transplant patient, and one normal individual (**Fig. 4a**). The presence of this mutation in a post-transplant patient and in a normal control suggests that *FOXO1^S22W^* can also arise prior to ALD or MASLD development. Consistent with this interpretation, we also identified *FOXO1^S22W^* in the liver tissue of a patient who received a second liver transplant. We also detected somatic mutations in *CIDEB* and *GPAM* (**Fig. 4b,c**), two genes with liver-enriched expression patterns whose mutations have primarily been reported in liver tissue^82–85^. Together, these findings strongly suggest that these somatic mutations, detected in cfDNA, originate from the liver.

**Fig. 4:**
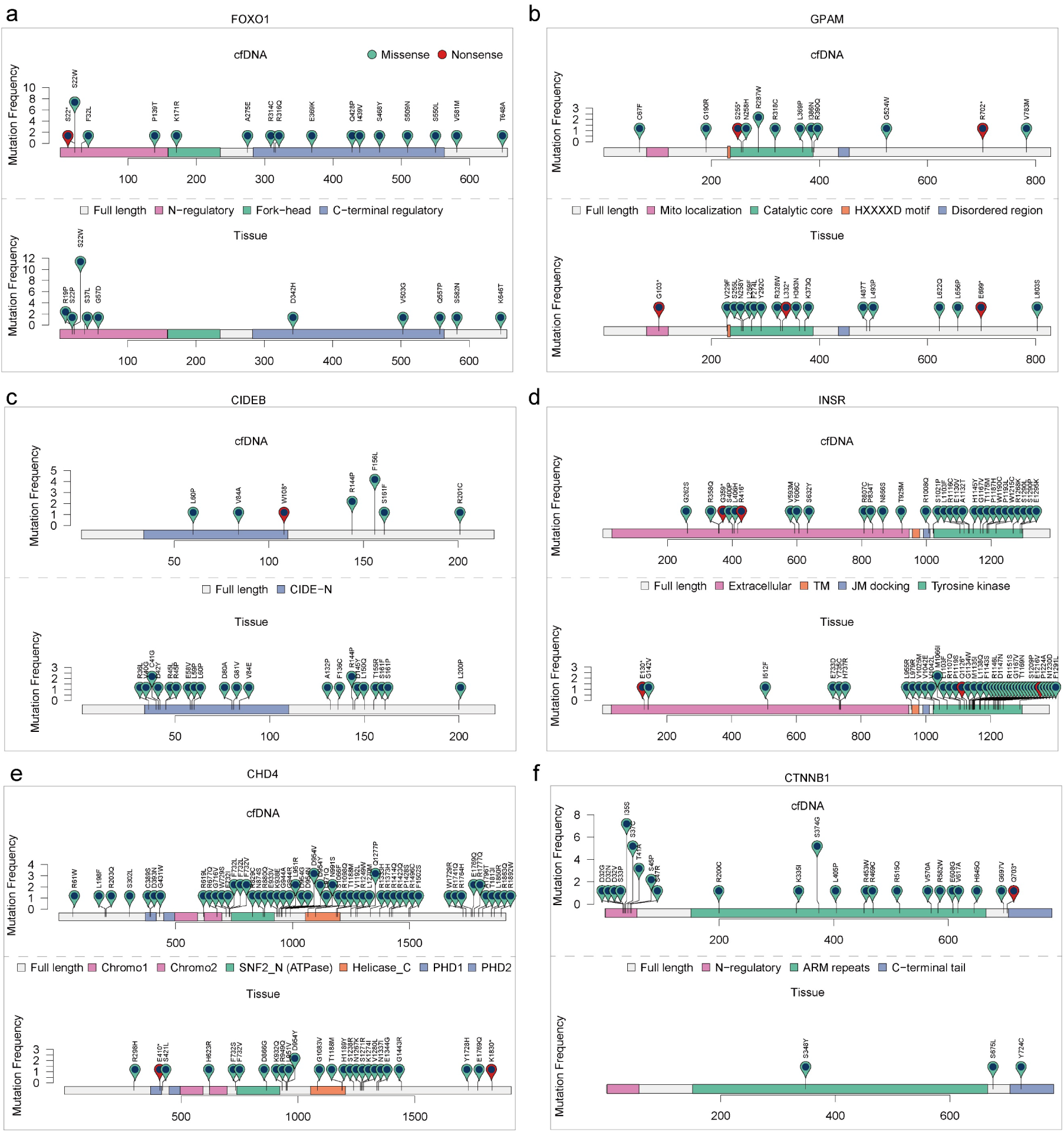
Mutation spectra of recurrently mutated genes in cfDNA and liver tissues. a–f. Lollipop plots showing the distribution of somatic mutations in *FOXO1* (a), *GPAM* (b), *CIDEB* (c), *INSR* (d), *CHD4* (e), and *CTNNB1* (f), the upper plot shows mutations detected in cfDNA, and the lower plot shows mutations detected in tissue. The x axis indicates amino acid position in the encoded protein, and the y axis indicates mutation frequency. Each circle represents a protein-altering mutation located at the corresponding amino acid position. Colors denote mutation class, with teal indicating missense mutations and red indicating nonsense mutations. The corresponding amino acid changes are labeled. Recurrent hotspot mutations are evident in multiple genes, including FOXO1 S22W.

We next compared mutation patterns observed in liver tissue (n = 31 samples from 31 patients) and cfDNA (n = 272 samples from 224 patients). For *FOXO1*, the recurrent S22W hotspot mutation was detected in both liver tissues and cfDNA (**Fig. 4a**). In liver tissue, 11 *FOXO1^S22W^* mutations were identified, one of which was also detected in the matched cfDNA sample from the same patient (**Extended Data Fig. 4a**). Similar concordance at the level of recurrent mutation patterns was observed for *GPAM*, *INSR*, and *CHD4* (**Fig. 4b–e**), which showed broadly consistent mutation spectra in tissue and cfDNA. These results suggest that cfDNA captures key features of liver tissue driver-gene mutation patterns.

In contrast, *CTNNB1* mutations were frequently detected in cfDNA from both normal controls and patients with liver disease, with multiple recurrent hotspot events observed across groups. This suggests that *CTNNB1* alterations detected in cfDNA should be interpreted cautiously, as they may not be exclusively liver-derived and could reflect contributions from non-hepatic cell populations or multiple tissues (**Fig. 4f, Extended Data Fig. 3d,e**). *CTNNB1* mutations were more frequently detected in female patients than in male patients (22/102 versus 15/103 patients). The mutational spectrum was dominated by canonical β-catenin regulatory hotspot mutations affecting residues within the N-terminal degradation motif, including D32, S33, S37, T41, and S45. Similar hotspot mutations were observed in both sexes (**Extended Data Fig. 4b**). Collectively, these findings indicate that while many driver mutations detected in cfDNA likely reflect liver-derived somatic alterations and recapitulate tissue-level mutational patterns, this may not be true for all genes.

### Distinct liver disease etiologies reveal specific mutational drivers

Different forms of CLD arise under distinct environmental and biological pressures, including alcohol exposure, overnutrition, and chronic viral infections from HBV or HCV^86–88^. Whether these etiologic contexts are associated with distinct somatic mutational preferences remains uncertain. We therefore compared cfDNA features across the major etiologies of liver disease. Among patients with cirrhosis, cfDNA abundance was significantly elevated relative to normal controls irrespective of etiology (**Fig. 5a**). Notably, cfDNA levels were higher in ALD than in MASLD matched for F4 cirrhosis, with median concentrations of 21.7 and 13.8 ng/μl, corresponding to increases of 153.8% and 61.4%, respectively, relative to normal controls. These findings suggest that the contribution of diseased tissue to the circulating cfDNA pool differs across etiologies. In contrast, analysis of mutation burden in CASMs genes revealed the opposite pattern: patients with MASLD showed a higher burden of mutations than patients with ALD (**Fig. 5b**). No such etiology-associated differences were observed for CH genes or non-disease control genes (**Extended Data Fig. 5a,b**), supporting the specificity of these differences with regard to CASM genes.

**Fig. 5:**
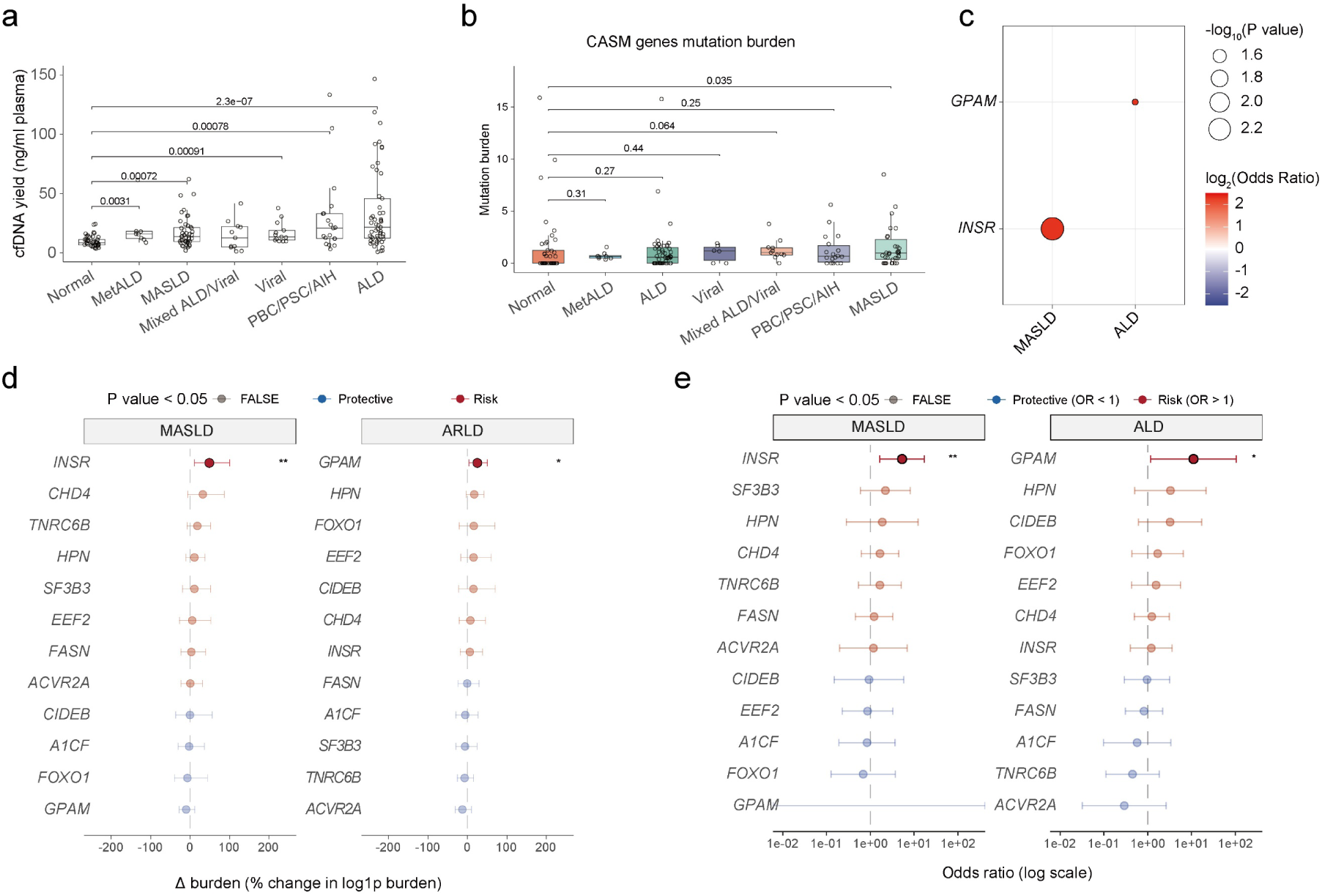
cfDNA yield, cirrhosis driver gene mutation burden, and gene-level etiology-associated analyses across disease groups. a. Comparison of plasma cfDNA yield across etiological groups. The y axis shows cfDNA yield (ng/ml plasma), and the x axis shows the Normal, ALD, MASLD, MetALD, Viral, Mixed ALD/Viral and PBC/PSC/AIH groups. P values for group comparisons are indicated. b. Comparison of mutation burden in CASM genes across etiological groups. The y axis shows mutation burden, and the x axis shows the Normal, ALD, MASLD, MetALD, Viral, Mixed ALD/Viral and PBC/PSC/AIH groups. P values for group comparisons are indicated. c. Summary of gene–etiology associations based on mutation presence analysis. The x axis shows etiological groups (MASLD and ALD), and the y axis shows genes. Bubble color indicates log2(odds ratio), and bubble size indicates -log10(P value). The plot summarizes the direction and significance of gene enrichment in specific etiological backgrounds. d. Gene-level etiology comparison based on mutation burden analysis. For the MASLD and ALD groups, the plot shows the change in burden for each gene relative to all other groups. The x axis indicates Δ burden (percent change in log1p-transformed burden). Point estimates and error bars represent the effect size and its interval. Red indicates the risk direction, and blue indicates the protective direction. Asterisks denote statistical significance (*P ≤ 0.05; **P ≤ 0.01). e. Gene-level etiology comparison based on mutation presence analysis. For the MASLD and ALD groups, the plot shows logistic regression results for the probability of mutation in each gene. The x axis shows the odds ratio (log scale). Point estimates and error bars represent the odds ratio and its confidence interval. Red indicates the risk direction (OR > 1), and blue indicates the protective direction (OR < 1). Asterisks denote statistical significance (*P ≤ 0.05; **P ≤ 0.01).

To further define mutational selection at the gene level, we fit covariate-adjusted regression models in F4 patients to estimate associations between individual disease etiologies and gene-specific mutation burden. These models adjusted for age, sex, race and sequencing read count. This analysis revealed distinct mutational patterns across etiologies: mutations in *INSR* were enriched in MASLD (P=0.009; FDR=0.11), whereas mutations in *GPAM* were enriched in ALD (P=0.022; FDR=0.26) (**Fig. 5d**). Analysis based on mutation presence yielded similar results (**Fig. 5c,e**). Notably, these cfDNA-derived patterns were concordant with our tissue-based dN/dS results, in which *INSR* showed stronger positive selection in MASLD, whereas *GPAM* showed stronger positive selection in ALD (**Fig. 1f,g, Extended Data Fig. 1c,d**).

To assess whether these associations were also observable across a broader disease spectrum, we performed an exploratory analysis including F0–F4 samples. Because these analyses were not adjusted for age, sex, or fibrosis stage, we interpreted them descriptively and used them to evaluate concordance in directional effect. In this exploratory analysis, MASLD showed concordant enrichment patterns, most notably for *INSR* and *CHD4* (**Extended Data Fig. 5c,d**). *INSR* showed the clearest signal, with a mean mutation burden approximately 3-fold higher in MASLD than in combined non-MASLD etiologies (permutation test, P = 0.0041; FDR = 0.053). *INSR* mutation presence was also higher in MASLD than in combined non-MASLD etiologies (23.2% versus 7.9%; P = 0.011; FDR = 0.14), and *INSR* mutation burden showed the same direction of effect when MASLD was compared with normal liver controls (permutation test, P = 0.027; FDR = 0.35). *CHD4* mutation burden showed a similar pattern, with an approximately 5-fold enrichment in MASLD relative to combined non-MASLD etiologies (permutation test, P = 0.046; FDR = 0.29) and a 7.0-fold enrichment relative to normal liver controls (permutation test, P = 0.072; FDR = 0.46). Consistent with this direction of effect, *CHD4* mutations were detected in 27.9% of MASLD cases, compared with 15.9% of combined non-MASLD etiologies (P = 0.079; FDR = 0.51) and 12.2% of normal liver controls (P = 0.10; FDR = 0.67). In the same exploratory F0–F4 analysis, *GPAM* showed a directionally consistent pattern, with higher mutation burden in ALD than in other disease etiologies (**Extended Data Fig. 5e,f**). Mean *GPAM* mutation burden was 3-fold higher in ALD than in combined non-ALD etiologies (permutation test, P = 0.009; FDR = 0.12). Consistent with this direction of effect, *GPAM* mutation presence was also higher in ALD than in combined non-ALD etiologies (13.2% versus 3.0%; Fisher’s exact test, P = 0.0098; FDR = 0.13). *GPAM* mutations were not detected in normal liver controls, whereas they were present in 13.2% of ALD cases (P = 0.017; FDR = 0.22). These exploratory results were directionally consistent with the covariate-adjusted F4 analysis and tissue-based dN/dS analysis (**Fig. 1f,g** and **Extended Data Fig. 1c,d**).

We next examined whether cfDNA fragmentation patterns differed across liver F4 cirrhosis etiologies. Relative to MASLD, ALD showed significantly lower contribution of the DNASE1L3-associated F-profile I, whereas MASLD showed a higher contribution of the DFFB-associated F-profile III compared with ALD (**Extended Data Fig. 5g,h**). These etiology-specific differences suggest that CLD subtypes differ not only in mutation spectra but also in the nuclease activities and/or cell-death programs that shape circulating cfDNA.

CfDNA analysis can capture the underlying disease-specific mutational landscape of the liver. The recurrent enrichment of *INSR* and *GPAM* across comparisons supports the notion that positively selected mutations in diseased liver tissue are shed into the circulation. The strongest signals were observed when MASLD/ALD was compared with normal controls or compared with all non-MASLD/non-ALD etiologies combined, whereas effect sizes were attenuated in individual pairwise comparisons, likely reflecting reduced statistical power in smaller subgroups. Although many associations did not remain statistically significant after correction for multiple testing, the consistent directionality of effects across comparisons supports their biological relevance. These findings suggest that distinct liver disease etiologies are associated with different programs of somatic mutation accumulation and/or clonal selection.

## Discussion

A key conceptual advance of this study is that cfDNA can be used to identify somatic mutations from non-malignant tissues. Although cfDNA-based mutation profiling has transformed oncology and prenatal diagnostics, its use in other organ disease setting has remained limited by the expectation that tissue-derived mutant fragments would be extremely rare in circulation. Using ultra-deep targeted duplex sequencing within a disease-focused driver gene framework, we show that liver-derived somatic signals can be detected in patients with CLD. For the liver, cfDNA captures key features of somatic genomics, including disease-associated driver mutations and etiology-specific mutational patterns. In this way, cfDNA provides access to mutational features that would otherwise require a liver biopsy. Across cfDNA and matched tissue analyses, CLD left a detectable mutational and fragmentation imprint in plasma. Taken together, these findings establish cfDNA as a minimally invasive approach for interrogating somatic evolution beyond cancer and suggest that this could be applicable to studying somatic evolution in other organs.

The transplantation analyses provide strong evidence for the hepatic origin of the mutations detected in plasma. We confirmed that a measurable fraction of circulating cfDNA after transplantation remained liver-derived. Moreover, transplantation was accompanied by a marked reduction in total cfDNA abundance and in mutation burden across CASM genes, whereas mutation burden in CH and non-disease control genes remained stable. These findings indicate that the disease-associated mutational signal detected in plasma originates predominantly from the diseased native liver rather than representing a nonspecific feature of circulating cfDNA. This transplantation-based design therefore provides a useful framework for linking plasma mutations to tissue of origin in non-malignant disease. Our results further suggest that different CLD etiologies are associated with distinct landscapes of somatic mutations. In both liver tissue and cfDNA, MASLD and ALD showed enrichment for different CASMs, consistent with etiology-specific patterns of somatic selection. These observations support a model in which different disease environments, including metabolic stress, alcohol exposure and viral injury, create distinct selective pressures that favor different hepatocyte clones. More broadly, this raises the possibility that somatic mutation patterns may provide a molecular readout of the biological forces shaping CLD.

This study has several limitations. First, although our cohort spans multiple liver disease etiologies, larger studies will be needed to refine the full spectrum of positively and negatively selected genes across CLD. Second, although transplantation provides strong evidence for liver origin, plasma cfDNA remains a composite signal contributed by multiple tissues, which may complicate interpretation for genes recurrently mutated outside the liver. Third, because our study used a targeted panel, our ability to detect etiology-specific selection may have been limited where relevant genes were not represented, particularly in viral and autoimmune liver disease. Despite these limitations, our study suggests that cfDNA can serve as a window into the molecular evolution of non-malignant tissues under chronic stress. CfDNA-based approaches may enable longitudinal tracking of clonal dynamics and molecular stratification of patients with CLD. In this way, liquid biopsy may extend beyond cancer as a strategy for studying and monitoring somatic evolution in chronic disease.

## Supporting information

Clinical characteristics of the study participants

dN/dS analysis across different sample groups

Sequencing statistics for study samples

Complete list of identified somatic mutations

## Acknowledgments

We acknowledge support from the Simmons Comprehensive Cancer Center (NCI Cancer Center Support Grant P30CA142543 and SCCC UTSW translational pilot grant), National Institutes of Health (DK139976).

## Author Contributions

BL, SR, APC, JL, TW, D.A.L performed bioinformatic analysis.

BL, MW, MZ performed cfDNA isolation, DNA isolation, and library prep. DA, AS, RW, CA performed statistical analysis.

DA, AS, RW, MP, AY, YH, AS, JT, LQ, and DH performed clinical analysis, obtained clinical samples, and provided scientific feedback.

BL, D.A.L, and HZ conceptualized and wrote the paper.

## METHODS

### Human samples

All human samples included in this study were collected with informed patient consent, in accordance with ethical guidelines and institutional regulations. UT-Southwestern Institutional Review Board (IRB) approval for the collection and analysis of tissue samples was granted under protocol #STU 062013-063, while blood sample collection was approved under protocol #STU 092013-010. In total, 303 clinical samples were analyzed. These included 31 liver tissue samples, and 272 plasma samples, including 163 pre-transplant plasma samples from patients with varying stages of liver disease, 53 post-transplant plasma samples, 41 healthy controls plasma samples and 15 heart disease plasma samples. The samples were collected and processed uniformly as described below, minimizing pre-analytical variability. Detailed clinical and demographic information was recorded for each sample, including patient age, gender, fibrosis stage, and disease etiology, as summarized in **Supplementary Table 1**.

### Plasma collection

Blood samples were drawn into Streck tubes, which are specifically designed to stabilize cfDNA. The samples were gently inverted several times to ensure proper mixing with the preservative agents. To isolate plasma, blood tubes were centrifuged at 3000 rpm for 10 minutes at 4°C. The supernatant plasma was carefully transferred to clean RNase/DNase-free tubes, taking care to avoid disturbing the buffy coat components. Plasma aliquots were immediately stored at -80°C until further processing.

### cfDNA extraction

cfDNA was extracted from plasma using the QIAamp Circulating Nucleic Acid Kit (Qiagen), following the manufacturer’s protocol. The DNA concentration was measured using the Qubit Fluorometer (ThermoFisher Scientific) with the Qubit dsDNA HS Assay Kit, which ensured precise quantification of low-concentration cfDNA. Quality assessment was performed using the Agilent TapeStation System with the High Sensitivity D5000 ScreenTape assay, verifying the expected cfDNA fragment size distribution and confirming the absence of high molecular weight genomic DNA contamination. Extracted cfDNA was stored at -80°C until library preparation.

### Liver tissue collection and DNA extraction

Liver tissue samples were collected from patients undergoing liver transplantation. Fresh tissue was placed into 50 mL conical tubes containing 20 mL of DMEM with 10% DMSO and kept on ice during transport to the laboratory. Upon arrival, the culture medium was discarded, and the tissue was immediately stored at -80°C until further processing. For DNA extraction, tissue samples were thawed on ice, and approximately 30 mg of tissue was sliced into 1–3 mm thick sections using a clean scalpel blade. The tissue slices were transferred to a 1.5 mL microcentrifuge tube and kept on dry ice. An adjacent section of 2–3 mm thickness was placed in a labeled cassette for formalin-fixation, paraffin-embedding (FFPE), and subsequent H&E staining. To avoid cross-contamination between samples, the scalpel blade was changed after each sample preparation. Genomic DNA was extracted from liver tissues using the protocol outlined by Samuel et al. in *Nature Protocols* (2013)^89^. To ensure high DNA integrity, tissues were manually homogenized with a pestle, followed by DNA extraction using the QIAshredder and QIAGEN AllPrep DNA/RNA Mini Kit. Extracted genomic DNA was stored at -80°C until further analysis.

### Library preparation and sequencing

#### Pre-capture library preparation

Sequencing libraries were prepared using the xGen™ cfDNA & FFPE DNA Library Prep v2 MC Kit (IDT) according to the manufacturer’s instructions. This generated dual-end UMI-labeled DNA fragments, enabling error correction and unique molecule tracking. Following library construction, the DNA concentration was quantified using the Qubit Fluorometer (ThermoFisher Scientific) with the Qubit dsDNA HS Assay Kit. Library quality and fragment size distributions were assessed using the Agilent TapeStation System.

#### Hybrid-capture library preparation

According to the manufacturer’s protocol of xGen™ hybridization capture of DNA libraries, 500 ng of each pre-capture library was pooled, combining 12 libraries into a single tube for hybridization. Target enrichment was performed using the xGen™ Custom Hyb Panel-Accel probes, which included a total of 7005 probes covering a genomic region of 201,496 bp. Library hybridization and capture were conducted using the xGen™ Hybridization Capture of DNA Libraries Kit (IDT), following the manufacturer’s protocol. This step enriched target regions for sequencing while maintaining library complexity. Post-capture libraries were purified using AMPure XP beads and amplified through PCR before proceeding to sequencing.

### Sequencing

Next-generation sequencing (NGS) was performed by Admera Health using the Illumina NovaSeq platform. Libraries were sequenced in a paired-end 150 bp (2×150) format, generating high-depth coverage for variant detection.

## Data analysis

### Mapping

Paired-end FASTQ files generated on the NovaSeq platform were first converted to an unmapped BAM (uBAM) format using FastqToSam in Picard (v3.4.0 ; Broad Institute, https://broadinstitute.github.io/picard/). UMI sequences were extracted from each read using ExtractUmisFromBam in fgbio (v2.5.21; Fulcrum Genomics) and stored in the RX tag of the resulting uBAM file for downstream consensus read generation. The uBAM was then converted back to FASTQ format using Picard SamToFastq (v3.4.0). Adapter sequences were removed using Skewer (v0.2.2)^90^, and the resulting reads were aligned to the hg38 reference genome using the mem algorithm in BWA (v0.7.17-r1188)^91^. Because alignment does not preserve RX tags, the aligned BAM was merged with the original uBAM using Picard MergeBamAlignment, thereby retaining UMI information in the aligned reads. As the IDT xGen UMI design comprises 32 predefined 8-bp UMI sequences, CorrectUmis in fgbio (v2.5.21) was used to correct sequencing errors in RX tags. Reads originating from the same source molecule were grouped using GroupReadsByUmi, after which single-strand consensus reads (SSCs) and duplex consensus reads (DCs) were generated using CallMolecularConsensusReads and CallDuplexConsensusReads, respectively. Consensus reads were subsequently realigned to the reference genome, and MergeBamAlignment was used again to restore tag information to the aligned consensus BAM.

### Mutation calling

To maximize specificity for rare variant detection in plasma cfDNA, we used a dedicated bioinformatics pipeline developed in the Landau Laboratory (Weill Cornell Medicine) to improve the accuracy of low-frequency variant detection. The code can be found in https://github.com/SriRaj34/yosemite and https://github.com/SriRaj34/yosemite_indels

### DN/dS selection analysis

To evaluate signals of selection in the targeted sequencing genes, we performed dN/dS analysis of coding somatic mutations using the dndscv^45^ R package. This method applies a maximum-likelihood framework for somatic mutation data to estimate gene-level selection pressure by comparing the observed and expected numbers of nonsynonymous and synonymous mutations, thereby identifying genes under positive or negative selection. The analysis was restricted to a predefined set of 59 target genes, using coding single-nucleotide variants (SNVs) detected in the samples as input. Because this study was based on a targeted gene panel, we used the same parameter settings as Lawson et al.: mingenecovs = 0, maxcovs = 10, onesided = TRUE, use_indel_sites = FALSE, max_muts_per_gene_per_sample = Inf, and max_coding_muts_per_sample = Inf. Genes with significant qallsubs_cv or pallsubs_cv values were considered to show evidence of overall selection, whereas significant qpos_cv/ppos_cv and qneg_cv/pneg_cv values indicated positive or negative selection, respectively. To evaluate selection signals at different stringency levels, we used three thresholds: q < 0.05, q < 0.1, and p < 0.05. Here, the p values and q values were derived from dNdScv: p values were obtained using likelihood-ratio tests, and q values were calculated using the Benjamini–Hochberg multiple-testing correction^45^. For the dataset from Ng et al., 2021^37^, the same analysis was performed using the same 59 target genes and identical parameter settings to ensure comparability, and the corresponding results are shown in **Extended Data Fig. 1c,d**.

### Identification of donor-derived germline mutations

To obtain a more stable estimate of the fraction of post-transplant cfDNA derived from the donor liver, donor-derived germline mutations were identified on the basis of SSC reads. First, all mutation sites supported by SSC reads were collected separately from pre- and post-transplant cfDNA samples. To minimize interference from recipient-derived germline variants and pre-existing background signals, all sites with VAF > 0.001 in pre-transplant cfDNA were excluded. In addition, only sites with sequencing depth above the tertile cutoff of the corresponding depth distribution in both pre- and post-transplant cfDNA samples were retained. After this filtering step, the remaining sites were defined as the set of potential candidate sites for each sample, denoted Gp. To identify and remove donor-derived germline mutations in post-transplant plasma samples, we further developed a hierarchical filtering procedure based on the distribution of VAFs across candidate sites. For each post-transplant sample, all sites in Gp were ranked in descending order of VAF. The difference in VAF between each pair of adjacent ranked sites was then calculated and denoted as diff, defined as the VAF of the current site minus that of the next ranked site. For each sample, the mean of all diff values was calculated to characterize the overall separation between high-VAF sites and the remaining sites. Across all post-transplant samples, the mean diff value for each sample was collected, and the 0.3 quantile of these values was defined as diff03, which served as a cross-sample reference threshold. For a given post-transplant sample, its own mean diff value was compared with diff03, and the larger of the two was used as the initial cutoff, to avoid setting the threshold too low in samples with globally small diff values. Next, within the diff series of that sample, all positions with diff > cutoff × 1.5 were identified, and the smallest such rank was taken as the boundary separating candidate germline mutations from the remaining sites. All high-VAF sites ranked before this boundary were classified as candidate donor-derived germline mutations. The resulting calls were then manually reviewed in the context of the overall VAF distribution of the post-transplant sample, and samples or sites that were clearly inconsistent with a germline mutation pattern were removed. This yielded a higher-confidence set of donor-derived germline mutations, denoted Gt. For each individual sample, because most germline heterozygous variants are expected to have a theoretical VAF of approximately 0.5 in purely donor-derived DNA, the fraction of cfDNA derived from the donor liver was estimated as 2 × the median VAF of all sites in the donor-derived germline mutation set Gt for that sample. To avoid potential confounding from multi-organ transplantation, only patients who underwent isolated liver transplantation were included for estimation of the liver-derived cfDNA fraction (**Fig. 2a,b**). An example is shown in **Extended Data Fig. 2a,b**.

### Mutation burden and mutation presence analyses

To assess the relationships between clinical features, etiological groups, continuous variables, and somatic mutational characteristics, we performed mutation burden and mutation presence analyses. For mutation burden analysis, standardized mutation burden was calculated at the sample level for individual genes or predefined gene sets. For a single gene, mutation burden was defined as the total number of mutant SSCs harboring SNVs detected in that gene divided by the total number of sequenced SSCs covering that gene, multiplied by (10^6^). For gene sets, including CASM genes, CH genes, and non-disease control genes, mutation burden was defined as the total number of mutant SSCs across all genes in the set divided by the total number of sequenced SSCs covering the corresponding gene set, multiplied by (10^6^). In group comparisons and regression analyses, mutation burden was treated as the dependent variable, and etiological group or continuous clinical variables were treated as independent variables. Genes with mutations detected in fewer than 5% of all samples were excluded from mutation presence and mutation burden analyses because of insufficient statistical power. Because mutation burden showed a right-skewed distribution, a log1p transformation was applied before modeling. Multivariable models included age, sex, and race clinical covariates to control for potential confounding. For mutation presence analysis, each gene in each sample was encoded as a binary variable: samples with at least one detected somatic mutation in that gene were assigned a value of 1, and all others were assigned 0. Logistic regression was then used to evaluate the associations of clinical variables, etiological groups, and other covariates with the probability of mutation in each gene. Multivariable models were further adjusted for age, sex, race, and sequencing read count.

As for exploratory analyses, differences in mutation presence between etiology groups were assessed using Fishers exact test, while differences in mutation burden were evaluated using the permutation test (10,000 permutations), in which case–control labels were randomly permuted to generate the empirical null distribution of the Wilcoxon rank statistic. All analyses were adjusted for multiple comparisons across genes.

### End motif analysis

End-motif analysis was performed using FinaleToolkit (v 0.11.0)^92^. Overlaped target regions in the custom capture panel were merged, and 30 bp were trimmed from both ends of each merged interval to minimize boundary effects. Four-mer end motifs were extracted from duplex-consensus BAM files using the interval-end-motifs function with a minimum mapping quality of 30 and fragment lengths of 50–300 bp. Counts of all 256 possible 4-mer end motifs were aggregated across all target regions to generate a sample-level end-motif profile. Expected motif frequencies were calculated from the hg38 reference genome within the trimmed target regions. Observed motif frequencies were normalized using an observed-to-expected (O/E) approach and rescaled to sum to one, generating normalized end-motif frequencies for downstream analyses. Normalized end-motif frequencies were CLR-transformed prior to dimensionality reduction. UMAP visualization was generated using the uwot (v 0.2.4) package following standard preprocessing ^93^.

### End-motif F-profile decomposition

To estimate the contribution of fragmentation signatures, we adopted the six reference F-profiles reported by Zhou et al^94^. The observed motif frequency vector of each sample was modeled as a non-negative linear combination of the six reference F-profiles using non-negative least squares (NNLS) implemented in the R package nnls. Estimated coefficients were normalized to sum to 100%, representing the relative contribution of each F-profile.

## Data availability

Sequencing data generated in this study will be deposited in the European Genome-phenome Archive (EGA) under accession number EXXX.

## Code availability

The SNV and indel calling pipelines used in this study were developed by the Landau Laboratory (Weill Cornell Medicine) and are publicly available at: https://github.com/SriRaj34/yosemite and https://github.com/SriRaj34/yosemite_indels

---

**Extended Data Fig. 1:**
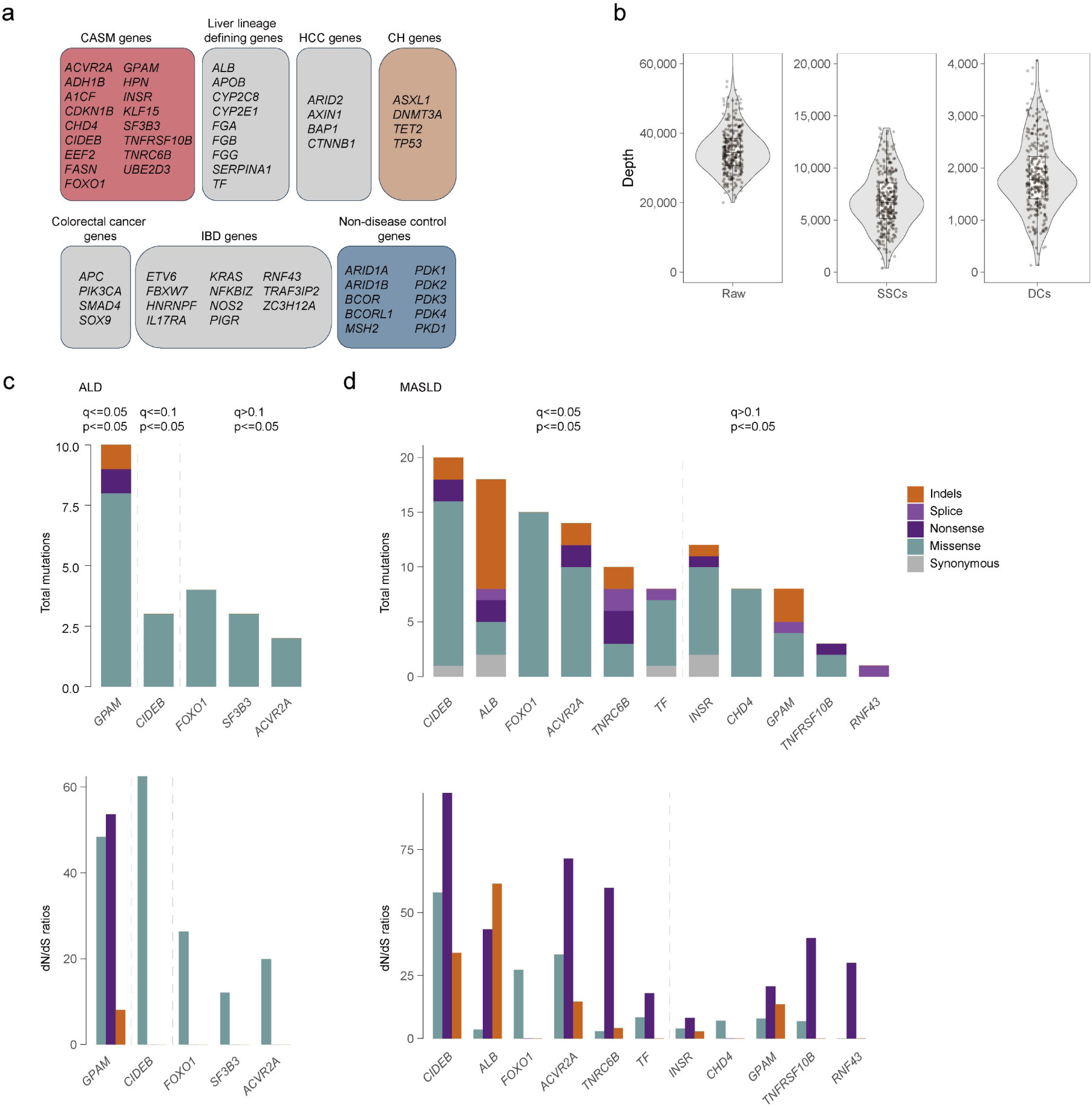
Sequencing coverage and selection analyses in different etiological subgroups. a. Classification of genes included in the targeted panel: Genes in the targeted panel were grouped into seven categories, including chronic liver disease (CLD)-associated somatic mutation (CASM) genes, liver lineage defining genes, hepatocellular carcinoma (HCC) genes, clonal hematopoiesis (CH) genes, colorectal cancer driver genes, inflammatory bowel disease (IBD) genes, and non-disease control genes. b. Distribution of sequencing coverage. Violin plots show the depth distributions for raw sequencing reads (Raw), single-strand consensus sequences (SSCs), and duplex consensus sequences (DCs). Each dot represents one sample, and boxplots indicate the median and interquartile range. c. Selection analysis in the ALD subgroup of the published dataset. The upper panel shows the total number of mutations detected in each gene, stratified by mutation class, and the lower panel shows the corresponding dN/dS ratios. Dashed lines indicate different statistical significance thresholds. P values and q values were derived from dNdScv. d. Selection analysis in the MASLD subgroup of the published dataset. The upper panel shows the total number of mutations detected in each gene, stratified by mutation class into synonymous, missense, nonsense, splice, and indel mutations; the lower panel shows the corresponding dN/dS ratios. Dashed lines indicate different statistical significance thresholds. P values and q values were derived from dNdScv.

**Extended Data Fig. 2:**
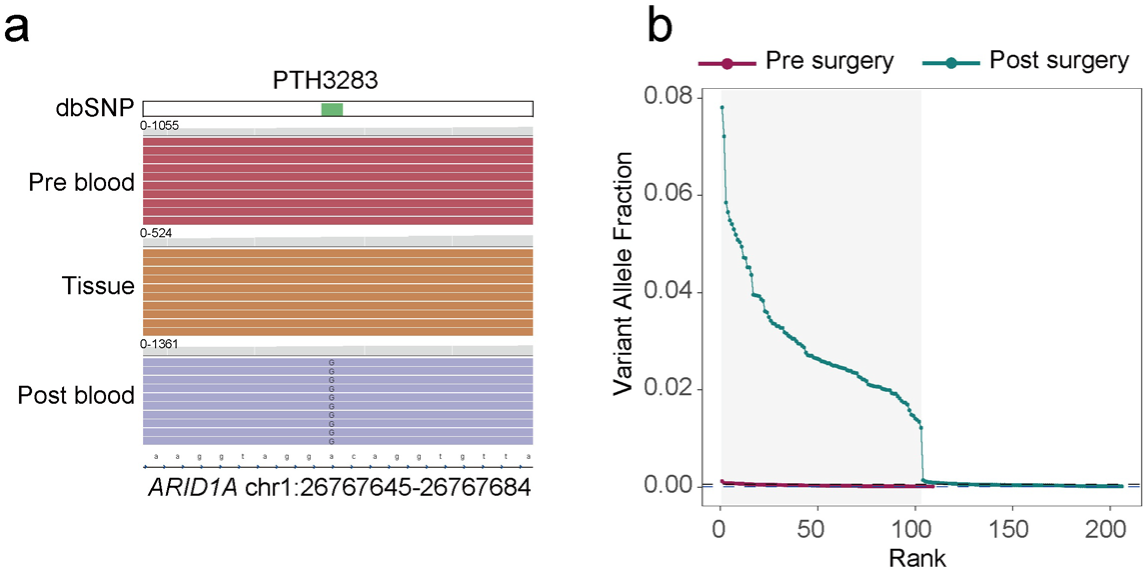
Detection of donor-derived cfDNA after liver transplantation using germline variants. a. Representative duplex sequencing reads at a donor-derived germline variant in *ARID1A* (PTH3283; chr1:26767645–26767684). The variant was absent in pre-transplant plasma (Pre blood) and recipient tissue (Tissue) but detected in post-transplant plasma (Post blood) at a variant allele fraction (VAF) of approximately 6%, consistent with the presence of donor-derived cfDNA following liver transplantation. b. Ranked variant allele fractions (VAFs) of candidate germline variants in post-transplant plasma samples, with matched pre-transplant plasma samples shown as controls. Variants were independently ranked by decreasing VAF in the post-transplant and pre-transplant samples. The shaded region denotes the putative germline variant set identified from the post-transplant samples for downstream donor-fraction estimation.

**Extended Data Fig. 3:**
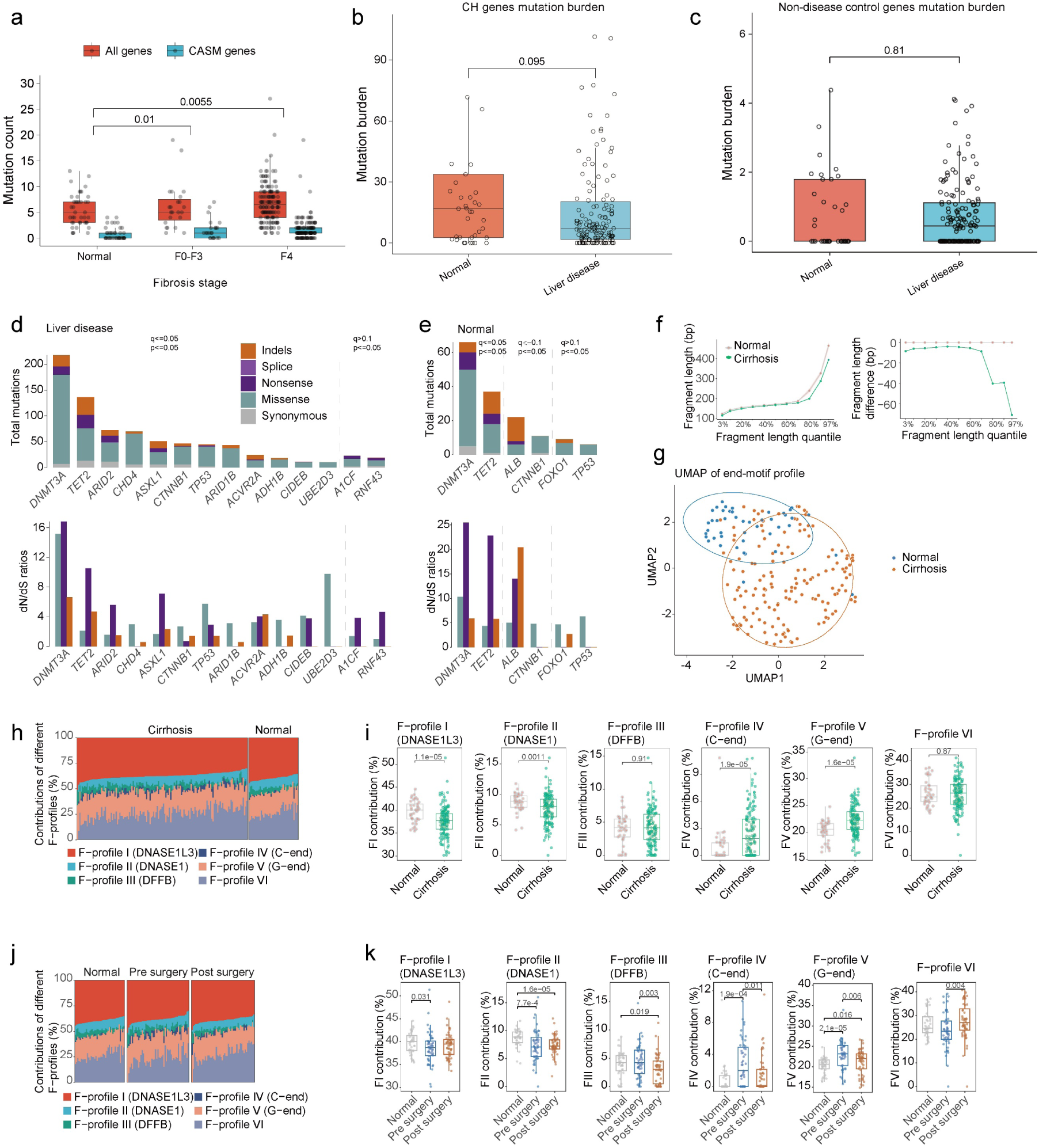
Mutation burden, selection signals, and fragmentomic features of cfDNA. a. Comparison of total mutation counts and mutation counts in CASM genes across fibrosis stage groups. Orange indicates mutations detected across the full gene panel, and blue indicates mutations in CASM genes. The x axis shows the Normal, F0–F3, and F4 groups, and the y axis shows mutation count. P values were calculated using binomial logistic regression to compare the fraction of mutations occurring in CASM genes among all detected coding mutations between groups. b. Comparison of mutation burden in CH genes across fibrosis stage groups. The x axis shows the Normal and liver disease (F0-F4) groups, and the y axis shows mutation burden. P values for group comparisons are indicated. c. Comparison of mutation burden in non-disease control genes across fibrosis stage groups. The x axis shows the Normal and liver disease (F0-F4) groups, and the y axis shows mutation burden. P values for group comparisons are indicated. d. Selection analysis in cfDNA samples from patients with liver disease. The upper panel shows the total number of mutations detected in each gene, and the lower panel shows the corresponding dN/dS ratios. Different colors denote different mutation classes, and dashed lines indicate different statistical significance thresholds. P values and q values were derived from dNdScv. e. Selection analysis in cfDNA samples from normal controls. The upper panel shows the total number of mutations detected in each gene, and the lower panel shows the corresponding dN/dS ratios. Dashed lines indicate different statistical significance thresholds. P values and q values were derived from dNdScv. f. Comparison of cfDNA fragment length distributions between normal controls and cirrhosis samples. The upper panel shows fragment lengths across fragment-length quantiles, and the lower panel shows fragment length differences relative to normal controls. The x axis indicates fragment-length quantiles, and the y axes indicate fragment length (bp) and fragment length difference (bp), respectively. g. UMAP analysis based on end-motif profiles. Each point represents one sample; blue indicates Normal and orange indicates Cirrhosis. The two groups show a tendency toward separation in low-dimensional space. h. Relative contributions of different F-profiles in cirrhosis and normal control samples. Each bar represents one sample, and the y axis shows the proportion (%) of each F-profile. F-profiles I–VI correspond to distinct nuclease-associated fragmentation features, including DNASE1L3-associated F-profile I, DNASE1-associated F-profile II, and DFFB-associated F-profile III. i. Comparison of the contributions of individual F-profiles between normal control and cirrhosis samples. Panels show the distributions of F-profile I (DNASE1L3), F-profile II (DNASE1), F-profile III (DFFB), F-profile IV (C-end), F-profile V (G-end), and F-profile VI. The y axis shows the corresponding F-profile contribution (%), and the x axis shows the Normal and Cirrhosis groups. P values for group comparisons are indicated. j. Stacked bar plots showing the relative contributions of six cfDNA fragmentation profiles (F1–F6) in normal controls, pre-surgery plasma, and post-surgery plasma. k. Quantification of the contribution of each fragmentation profile in normal controls, pre-surgery plasma, and post-surgery plasma. Boxes indicate median and interquartile range; whiskers denote 1.5 x the interquartile range; points represent individual samples; P values are shown for the indicated comparisons.

**Extended Data Fig. 4:**
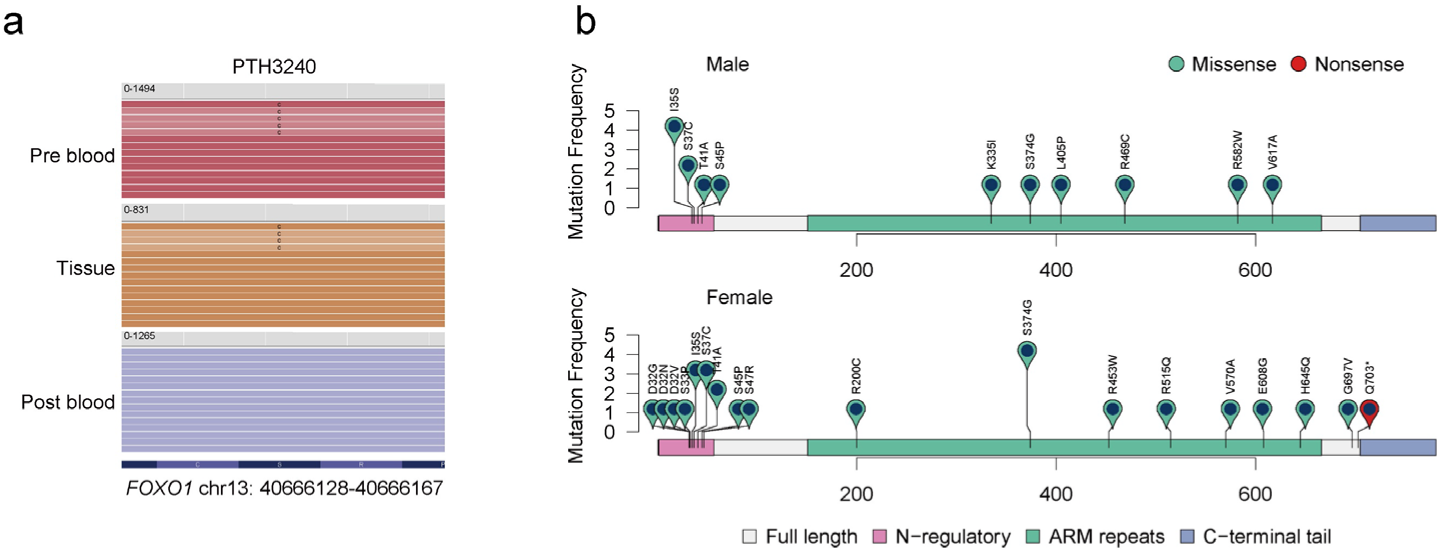
FOXO1 S22W mutation detected in pre-transplant cfDNA and liver tissue but not post-transplant cfDNA. a. Representative read alignments at the *FOXO1* locus (chr13:40666128–40666167) from patient PTH3240, showing detection of the S22W mutation in pre-transplant plasma cfDNA and matched liver tissue, but not in post-transplant plasma cfDNA. The mutant allele is marked by the substituted base within aligned reads. Numbers at the top of each track indicate total read depth across the displayed region. b. Lollipop plots showing the distribution of somatic mutations in *CTNNB1* detected in cfDNA from male and female patients. The x axis indicates amino acid position in the encoded protein, and the y axis indicates mutation frequency. Each circle represents a protein-altering mutation located at the corresponding amino acid position. Colors denote mutation class, with teal indicating missense mutations and red indicating nonsense mutations.

**Extended Data Fig. 5:**
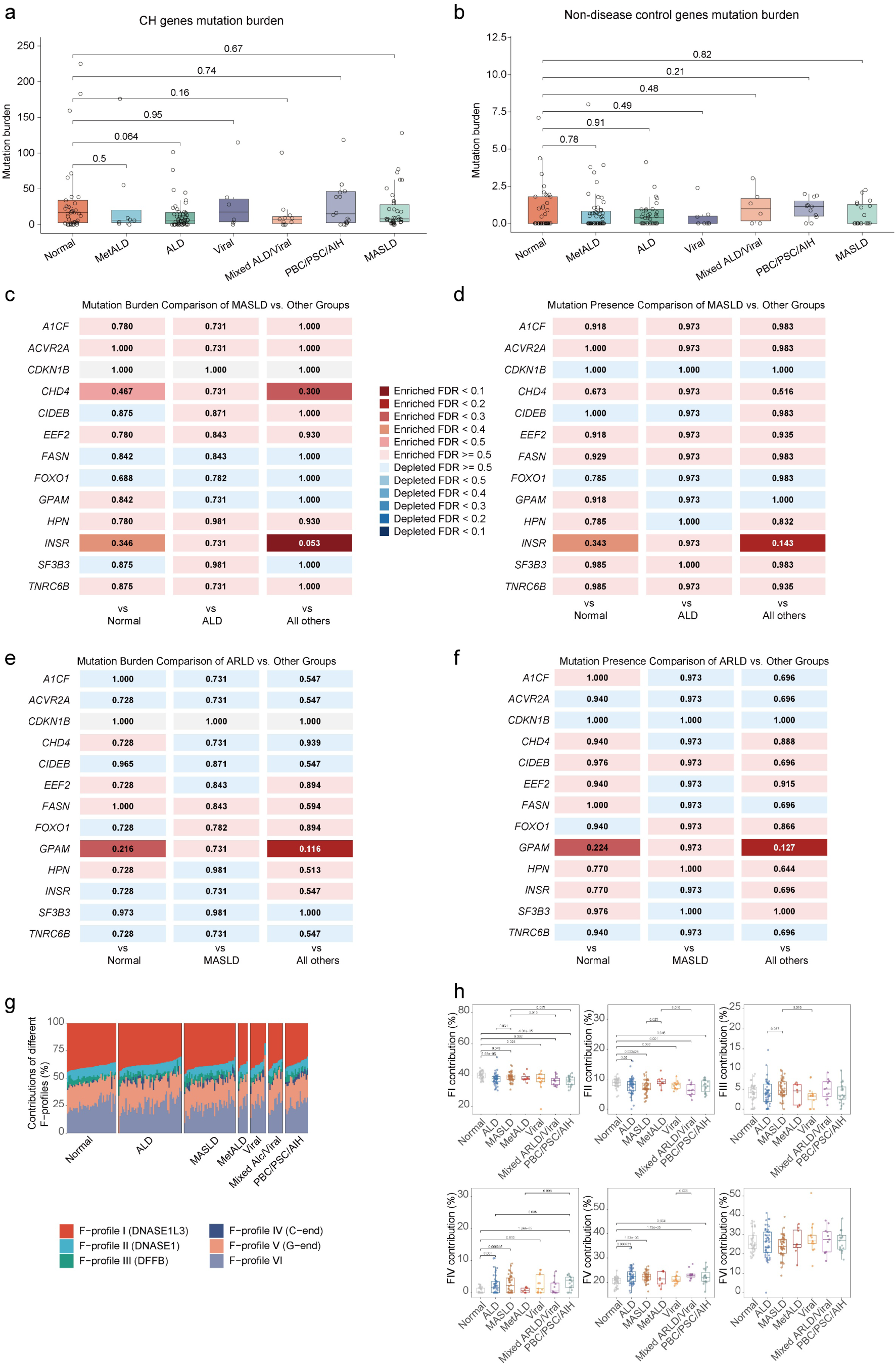
Etiology-associated patterns of cfDNA mutations and fragment-end features. a. Comparison of mutation burden in CH genes across etiological groups. The x axis shows etiological categories, and the y axis shows mutation burden. P values for group comparisons are indicated. b. Comparison of mutation burden in non-disease control genes across etiological groups. The x axis shows etiological categories, and the y axis shows mutation burden. P values for group comparisons are indicated. c-f. Heat maps summarizing etiology-stratified comparisons of gene-specific mutation burden and mutation presence across recurrent CASM genes detected in cfDNA. c, Comparison of mutation burden in MASLD versus normal controls, ALD, and all other etiologies combined. d, Comparison of mutation presence in MASLD versus normal controls, ALD, and all other etiologies combined. e, Comparison of mutation burden in ALD versus normal controls, MASLD, and all other etiologies combined. f, Comparison of mutation presence in ALD versus normal controls, MASLD, and all other etiologies combined. Numbers shown in each cell indicate FDR values for the corresponding comparison. Red indicates relative enrichment in the focal etiology, whereas blue indicates relative depletion; color intensity reflects the strength of the association after multiple-testing correction. g. Relative composition of cfDNA fragment-end features (F-profiles) across etiological groups. Each bar represents one sample, and the y axis shows the relative contribution (%) of each F-profile. Colors denote F-profile I (DNASE1L3), F-profile II (DNASE1), F-profile III (DFFB), F-profile IV (C-end), F-profile V (G-end), and F-profile VI. Samples are grouped by etiology, including Normal, ALD, MASLD, MetALD, Viral, Mixed ALD/Viral, and PBC/PSC/AIH. h. Comparison of the relative contributions of individual F-profiles across etiological groups. The six boxplots correspond to F-profiles I–VI. The y axis shows the corresponding F-profile contribution (%), and the x axis shows etiological categories. P values for group comparisons are indicated.

